# Spatial and temporal diversity of astrocyte phenotypes in Spinocerebellar ataxia type 1 mice

**DOI:** 10.1101/2021.09.13.460129

**Authors:** Juao-Guilherme Rosa, Katherine Hamel, Carrie Sheeler, Ella Borgenheimer, Stephen Gilliat, Alyssa Soles, Fares Ghannoum, Kaelin Sbrocco, Hillary P. Handler, Orion Rainwater, Ryan Kang, Marija Cvetanovic

## Abstract

While astrocyte heterogeneity is an important feature of the healthy brain, less is understood about spatiotemporal heterogeneity of astrocytes in brain disease. Spinocerebellar ataxia type 1 (SCA1) is a progressive neurodegenerative disease caused by a CAG repeat expansion mutation in the gene *Ataxin1* (*ATXN1*). We characterized astrocytes across disease progression in the four clinically relevant brain regions, cerebellum, brainstem, hippocampus, and motor cortex of *Atxn1*^*154Q/2Q*^ mice, a knock-in mouse model of SCA1. We found brain region specific changes in astrocyte density, GFAP expression and area, early in disease and prior to neuronal loss. Expression of astrocytic core homeostatic genes was also altered in a brain-region specific manner and correlated with neuronal activity indicating that astrocytes may compensate or exacerbate neuronal dysfunction in a brain region specific manner. Late in disease, expression of astrocytic homeostatic genes was reduced in all four brain regions indicating loss of astrocyte functions. We observed spatiotemporal changes in microglia with no obvious correlation with spatiotemporal astrocyte alterations indicating a complex orchestration of glial phenotypes in disease. These results support spatiotemporal diversity of glial phenotypes as an important feature of the brain disease that may contribute to SCA1 pathogenesis in a brain-region and disease stage-specific manner.

## 1. Introduction

Astrocytes play critical roles in brain development and in functioning of the adult brain [1][2][3]. One key way that astrocytes regulate neuronal activity is by maintaining homeostasis of ions, neurotransmitters and water in the brain [4][5][6][7]. Differences in the activity and metabolic needs of neurons across and within brain regions indicate the heterogeneous and region-specific activity of astrocytes [8][9][10][11][12][13][14]. While heterogeneity of astrocyte morphology was described by Ramon Y Cajal, recent advances in single-cell RNA sequencing (scRNA-seq), spatial transcriptomics, and single-cell mapping of astrocyte interactions have begun characterizing the molecular heterogeneity of astrocytes and its relevance for healthy brain function [15][16][17].

In brain diseases, astrocytes undergo phenotypic change termed reactive gliosis and defined as morphological and gene expression changes that may result in altered astrocyte function [18]. Reactive astrocytes contribute to pathogenesis of neurodegenerative disease and can either exacerbate or ameliorate neurodegeneration as has been shown in Huntington’s disease (HD) [20][21], Alzheimer’s disease (AD) [22][23][24] and Spinocerebellar ataxia type 1 (SCA1) [25]. How regional heterogeneity of astrocytes affects their changes in disease and whether it contributes to the selective regional neuronal vulnerability, one of the key features of neurodegenerative diseases [26], remain important unanswered questions [27].

SCA1 is an inherited neurodegenerative disorder caused by CAG repeat expansion mutations in the *Ataxin-1* (*ATXN1)* gene [28]. Repeat expansions of about 39 or more CAG repeats result in the expression of a toxic polyglutamine (polyQ) ATXN1 protein [29]. SCA1 is characterized by progressive degeneration that is most severe in the cerebellum and brainstem, but is also present in other brain regions, and manifests behaviorally as deficits in motor coordination (i.e. ataxia), swallowing, speaking, cognition, and mood [30][31][32]. While *Atxn1*-targeting antisense oligonucleotides (ASOs) show promise in pre-clinical trials [33], there are currently no disease-modifying treatments available for SCA1.

Motor deficits and underlying cerebellar pathology have been the focus of the majority of SCA1 studies so far. However, *ATXN1* is expressed throughout the brain and likely affects brain regions beyond the cerebellum [34][35]. Pathology is described in the brainstem, motor cortex, and hippocampus of SCA1 patients, and is thought to contribute to SCA1 symptoms such as cognitive deficits, mood disorders, difficulties in respiration and swallowing, and premature lethality [36][37][38][39][40][41]. We have previously demonstrated that astrocytes actively contribute to disease pathogenesis in the cerebellum of SCA1 mice [25], but how astrocytes are altered in other SCA1 affected brain regions remains unknown.

To better understand brain-wide SCA1 astrocyte pathology we have utilized the *Atxn1*^*154Q/2Q*^ mouse model of SCA1 wherein 154 CAG repeats have been knocked-in into the endogenous mouse *Atxn1* gene [42]. Because the *Atxn1* gene is expressed under the endogenous promoter, *Atxn1*^*154Q/2Q*^ mice express polyQ ATXN1 in cells throughout central nervous system [https://www.proteinatlas.org/] and in brain cells such as neurons, astrocytes, and microglia [http://www.brainrnaseq.org/]. As a result, in addition to motor deficits, the *Atxn1*^*154Q/2Q*^ mice exhibit a premature lethality, failure to gain weight, and cognitive deficits [42][43].Furthermore, there is evidence of hippocampal and brainstem changes in *Atxn1*^*154Q/2Q*^ mice, although pathology in these brain regions remains to be thoroughly characterized [42][44][45].

To investigate region-specific and temporal contribution of astrocytes to SCA1, we compared morphology and gene expression changes in astrocytes in the cerebellum, brainstem hippocampus, and motor cortex of 12-and 24-week-old *Atxn1*^*154Q/2Q*^ mice. We focused on these time points because they represent early to mid and late disease stages. Early stages of disease are particularly of interest due to previous studies showing the significant potential of early therapeutic approaches [46][47].

We found diverse brain region specific morphological and gene expression changes in astrocytes in SCA1 mice. Of importance, in the brainstem we found a significant decrease in astrocytic density and GFAP expression, and in the expression of core astrocytic homeostatic genes including *Kcnj10, Aqp4, Slc1a2* and *Gja1*. These astrocytic genes play a critical role in the maintenance of brain homeostasis by regulating local concentrations of potassium (*Kir4*.*1*), water (*Aqp4*), and glutamate (*Slc1a2*)[48][7][49][50]. These results indicate atrophy and loss of key functions of brainstem astrocytes that may exacerbate SCA1 pathogenesis in this region. On the other hand, we found increased expression of core homeostatic genes in the cortical astrocytes indicating potentially beneficial roles of astrocytes in the SCA1 affected cortex.

We also investigated microglial and neuronal changes across these four brain regions. We found increased microglial density both in the brainstem and cortex despite these regions showing very different changes in astrocytes, indicating a complex and brain region specific relationship between SCA1 astrocytes and microglia. Importantly, we did not observe any changes in neuronal numbers in any of the examined regions at this early disease stage, indicating that early in disease glial changes precede neuronal loss in this SCA1 mouse model.

Finally, we investigated how regional SCA1 astrocyte phenotypes change with disease progression. Late in diseases, we found a homogeneously reduced expression of astrocyte homeostatic genes in all four brain regions. These results indicate spatial and temporal diversity of astrocyte phenotypes in SCA1 and that disease progression leads to a potential brain wide loss of astrocytic functions.

## 2. Materials and Methods

### Mice

The creation of the *Atxn1*^*154Q/2Q*^ mice was previously described [42]. Because repeat length in trinucleotide repeat expansions is unstable and prone to further expansion [51], we routinely perform genetic sequencing of our mouse lines. We found that the number of repeats has recently expanded in our colony from 154 CAG to 166 CAG. Based on previous studies this increase in CAG repeat number is expected to increase severity of disease and lower age of symptom onset. We used approximately equal number of male and female mice in all experiments.

### Immunofluorescent (IF) staining

IF was performed on a minimum of six different floating 45-μm-thick brain slices from each mouse (six technical replicates per mouse per region or antibody of interest). Confocal images were acquired using a confocal microscope (Olympus FV1000) using a 20X or 40X oil objective. Z-stacks consisting of twenty non-overlapping 1-μm-thick slices were taken of each stained brain slice per brain region (i.e., six z-stacks per mouse, each taken from a different brain slice). The laser power and detector gain were standardized and fixed between mice within a surgery cohort, and all images for mice within a cohort were acquired in a single imaging session to allow for quantitative comparison.

We used primary antibodies against Purkinje cell marker calbindin (mouse, Sigma-Aldrich, C9848), neuronal marker neuronal nuclei (NeuN) (rabbit, Abcam, Ab104225), astrocytic marker glial fibrillary acidic protein (GFAP) (chicken, Millipore, AB5541), microglial marker ionized calcium binding adaptor molecule 1 (Iba1) (rabbit, WAKO, 019-19741), c-Fos, PSD-95 and vesicular glutamate transporter 2 (VGLUT2) (guinea pig, Millipore, AB2251-I) as previously described [25][52]. Quantitative analysis was performed using ImageJ (NIH) as described previously. To quantify relative intensity of staining for GFAP and calbindin we measured the average signal intensity in the region of interest and normalized it to that of the WT littermate controls. The density of neurons, astrocytes, microglia, and active neuronal cells was determined by normalizing the number of NeuN+, GFAP+, Iba1+, and c-Fos++ cells respectively with the area of the region of interest. We determined GFAP+ and Iba1+ percent area by creating a mask of GFAP and Iba1 staining respectively and recording the fraction of the region of interest covered by staining. To quantify atrophy of the cerebellar molecular layer we took six measurements per image of the distance from the base of the Purkinje soma to the end of their dendrites, the average being the molecular layer width for that image. Recession of climbing fibers was quantified by normalizing the width of VGLUT2 staining to the width of the molecular layer. The width of hippocampal neuronal layers (CA2, and CA3) was measured by dividing the area of each neuronal layer by the length of that layer as previously described [44].

### RNA Extraction, Sequencing and Analyses

Cerebellum, medulla, cerebral cortex, and hippocampal tissue was isolated from 26-week-old wild-type and *Atxn1*^*154Q/2Q*^ mice and stored in RNAlater solution (Thermo Fisher Scientific). Total RNA was isolated using TRIzol reagent (Thermo Fisher Scientific) following the manufacturer’s protocols. Tissue was homogenized using RNase-Free disposable pellet pestles in a motorized chuck. Purified RNA was sent to the University of Minnesota Genomics Center for quality control, including quantification using fluorimetry via RiboGreen assay kit (Thermo Fisher Scientific) and RNA integrity was assessed via capillary 34 electrophoresis using an Agilent BioAnalyzer 2100 to generate an RNA integrity number (RIN). RIN values for submitted RNA were above 8.0 for all samples except one medulla sample (RIN = 6.8). All submitted RNA samples had greater than 1µg total mass. Library creation was completed using oligo-dT purification of polyadenylated RNA, which was reverse transcribed to create cDNA. cDNA was fragmented, blunt ended, and ligated to barcode adaptors. Libraries were size selected to 320 bp ± 5% to produce average inserts of approximately 200 bp, and size distribution was validated using capillary electrophoresis and quantified using fluorimetry (PicoGreen, Thermo Fisher Scientific) and qPCR. Libraries were then normalized, pooled, and sequenced on an S4 flow cell by an Illumina NovaSeq 6000 using a 150-nucleotide, paired-end read strategy. The resulting FASTQ files were trimmed, aligned to the mouse reference genome (GRCm38), sorted, and counted using the Bulk RNAseq Analysis Pipeline from the Minnesota Supercomputing Institute’s Collection of Hierarchical UMII/RIS Pipelines (v0.2.0) (Baller et al., 2019). Genes less than 300 bp are too small to be accurately captured in standard RNAseq library preparations, so they were discarded from all downstream analyses.

Differential gene expression analysis was performed using the edgeR package (McCarthy et al., 2012; Robinson et al., 2009) (v3.30.3) in R (R Foundation for Statistical Computing v3.6.1). All four brain regions were analyzed independently. Genes with fewer than 10 counts across all samples in each region were excluded. Genes with FDR values less than or equal to 0.05 were considered significant.

### Reverse transcription and quantitative polymerase chain reaction (RT-qPCR)

Total RNA was extracted from dissected mouse cerebella, brainstem, cortex, and hippocampus using TRIzol (Life Technologies), and RT-qPCR was performed as described previously [8]. We used IDT Primetime primers (IDT). Relative mRNA levels were calculated using 18S RNA as a control and wild-type mice as a reference using 2^-ΔΔ-Ct^ as previously described [8].

### Data availability

All the data from this study are available at the request from the authors.

### Statistics

Wherever possible, sample sizes were calculated using power analyses based on the standard deviations from our previous studies, significance level of 5%, and power of 90%. Statistical tests were performed with GraphPad Prism 7.0. Data was analyzed using two-way ANOVA (to assess the impact of genotype and treatment) followed by post-hoc two-tailed, unpaired Student’s t-test (to test statistical significance of differences between groups of interest wherein only one variable was different (treatment or genotype), or one-way ANOVA followed by the Sidak’s post-hoc test. Outliers were determined using GraphPad PRISM’s Robust regression and Outlier removal (ROUT) with a Q=1% for non-biased selection.

## 3. Results

### Spatial diversity of astrocyte morphology during early disease stages in Atxn1^154Q/2Q^mice

Ataxin-1 is widely expressed throughout the brain in humans and in mice [53][54][44], yet how are astrocytes altered across relevant brain regions in SCA1 is unknown [35]. We first investigated morphological changes in astrocytes in the cerebellum, brainstem, hippocampus, and motor cortex of *Atxn1*^*154Q/2Q*^ mice during early disease stage. These regions have presumptive roles in motor deficits, premature lethality, and cognitive dysfunction in SCA1. For instance, aspiration pneumonia is one of the leading causes of death in patients with SCA1 [36]. Previously described neuronal degeneration and gliosis in the brainstem of patients with SCA1 are thought to contribute to the loss of ability to protect airways and, subsequently, aspiration pneumonia and death [36][55]

To investigate morphological changes in SCA1 astrocytes we quantified an increase in area occupied by glial fibrillary acidic protein (GFAP) (percentage of area that is GFAP+), a morphological characteristic of hypertrophy in reactive astrocytes, an increase in GFAP immunoreactivity, often used as a measure of reactive astrogliosis, and cell density where possible [56][57]. In the cerebellum, we examined Bergmann glia (BG), a subtype of cerebellar astrocytes that reside in the Purkinje and molecular layers and have a very intimate structural and functional relationship with Purkinje Cells (PCs), neurons affected in SCA1 [58][59]. We detected a significant increase in GFAP+ percent area (Figure 1A), and in the intensity of GFAP staining [60] indicative of hypertrophy of SCA1 Bergmann glia. We have previously described SCA1 alterations in the dentate gyrus of the hippocampus in *Atxn1*^*154Q/2Q*^ mice [45]. We examined astrocyte morphology and found trending increase in intensity of GFAP expression and GFAP+ astrocyte percent area, indicative of astrocyte hypertrophy in the dentate gyrus of *Atxn1*^*154Q/2Q*^ mice relative to wild-type controls (Figure 1B). In the brainstem, we characterized astrocyte morphology in the inferior olivary nucleus (ION), because of its role in cerebellar learning [41], because postmortem analysis of patient samples showing severe loss of volume in the ION [34], and due to high-quality of GFAP staining relative to other brainstem nuclei (Figure 1C). Interestingly, we found that in the inferior olive of *Atxn1*^*154Q/2Q*^ mice the intensity of GFAP staining and density of astrocytes were significantly reduced. Previous brain-wide investigation demonstrated neuronal loss in the primary motor cortex of SCA1 patients [34]. Because of the reported differences in astrocytes in gray and white matter, we assessed astrocyte morphology in layer 6 of the motor cortex and the underlying corpus callosum of *Atxn1*^*154Q/2Q*^ mice (Figure 1D). We detected a slight increase in GFAP intensity and percent area in the corpus callosum of *Atxn1*^*154Q/2Q*^ mice that did not reach statistical significance and may indicate emerging hypertrophy of cortical astrocytes (Figure 1D). There were no changes in astrocyte density, GFAP intensity or area in the layer 6 of motor cortex.

**Figure 1.**
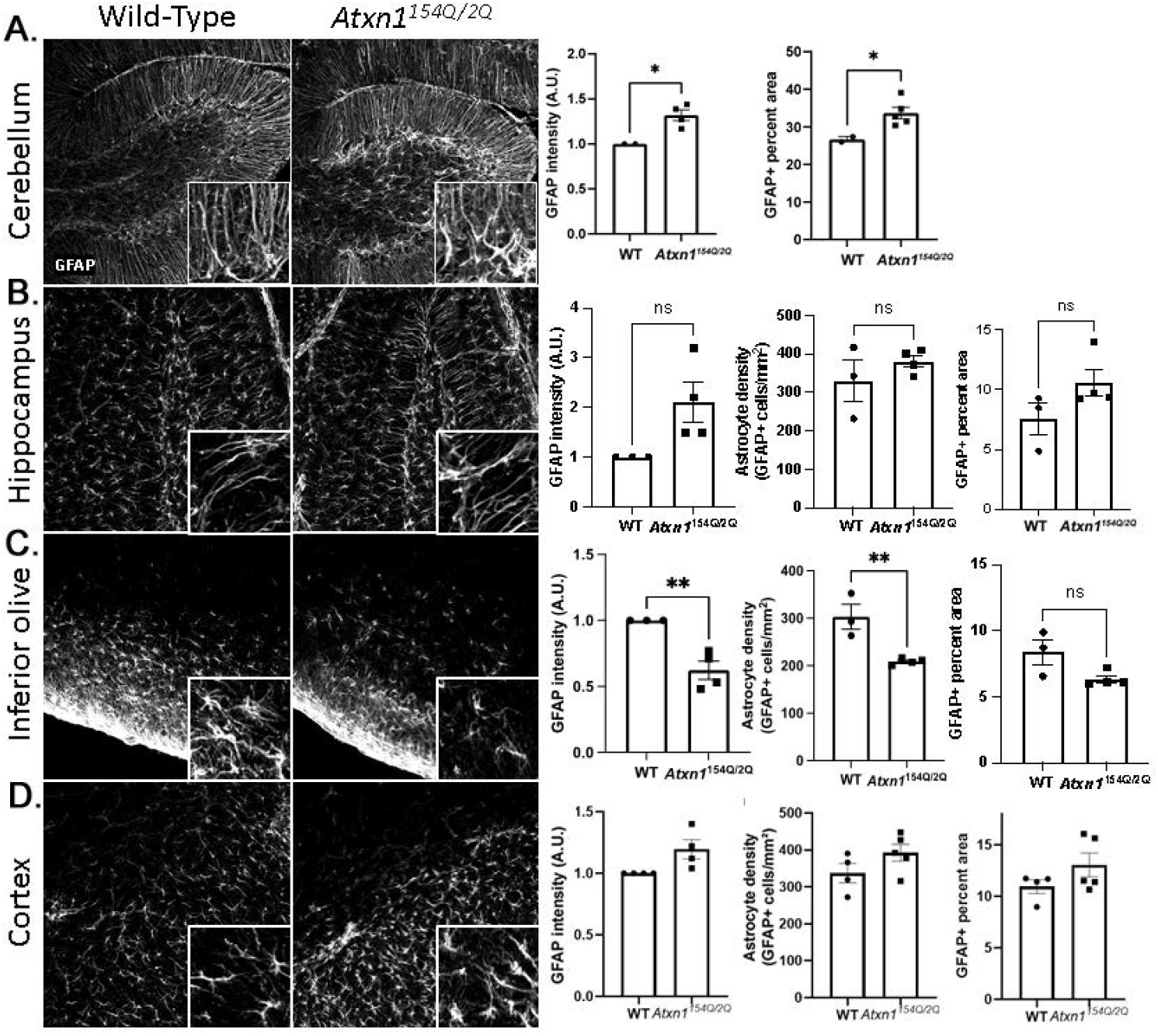
Spatial diversity of astrocytic morphology in *Atxn1*^*154Q/2Q*^ mice early in disease. Brain sections from 12 weeks old *Atxn1*^*154Q/2Q*^ and wild-type littermate controls (N = 3-5) were stained for GFAP. Confocal images of cerebellum (A, lobule X), hippocampus (B, dentate gyrus), brainstem (C, olivary nucleus), and motor cortex (layer 6 and corpus callosum) were used to quantify GFAP intensity, density of astrocytes and GFAP area. Data is presented as mean ± SEM with average values for each mouse represented by a dot. * p<0.05 Students t-test.

These results indicate diverse morphological changes in SCA1 astrocytes across brain regions with astrocytic loss and atrophy in the brainstem, astrocyte hypertrophy in the cerebellum and trending, but not significant astrocytic hypertrophy in the hippocampus and corpus callosum.

### Spatial diversity of astrocytic gene expression changes in SCA1

We next investigated molecular changes in SCA1 astrocytes across brain regions. Among the astrocytic roles critical for neuronal function is maintaining the homeostasis of extracellular environment. Astrocyte accomplish this by removing excess extracellular glutamate via glutamate transporters encoded by *Slc1a3* and *Slc1a2* genes, potassium ions via potassium channel encoded by *Kcnj10*, and water through the expression of Aquaporin-4, encoded by *Aqp4*)[61][62][63][64]. Astrocyte intercommunication in a network allowing for calcium wave propagation, potassium and glutamate buffering is facilitated by Connexin43 encoded by *Gja1*. We compared the molecular changes in SCA1 astrocytes across the cerebellum, brainstem, hippocampus, and cortex by quantifying the expression of these critical astrocytic core genes using RT-qPCR.

We found a significant reduction in the expression of *Aqp4, Gja1, Slc1a2*, and *Kcnj10*, in the brainstem (Figure 2A) and hippocampus (Figure 2B) of *Atxn1*^*154Q/2Q*^ mice at 12 weeks. In contrast, expression of these genes was increased in the cortex (Figure 2C). Given the importance of these homeostatic astrocytic genes for neuronal function, our results may indicate a novel mechanism by which astrocytes exacerbate neuronal dysfunction in the brainstem and hippocampus and compensate for or ameliorate dysfunction in the cortex of *Atxn1*^*154Q/2Q*^ mice.

**Figure 2.**
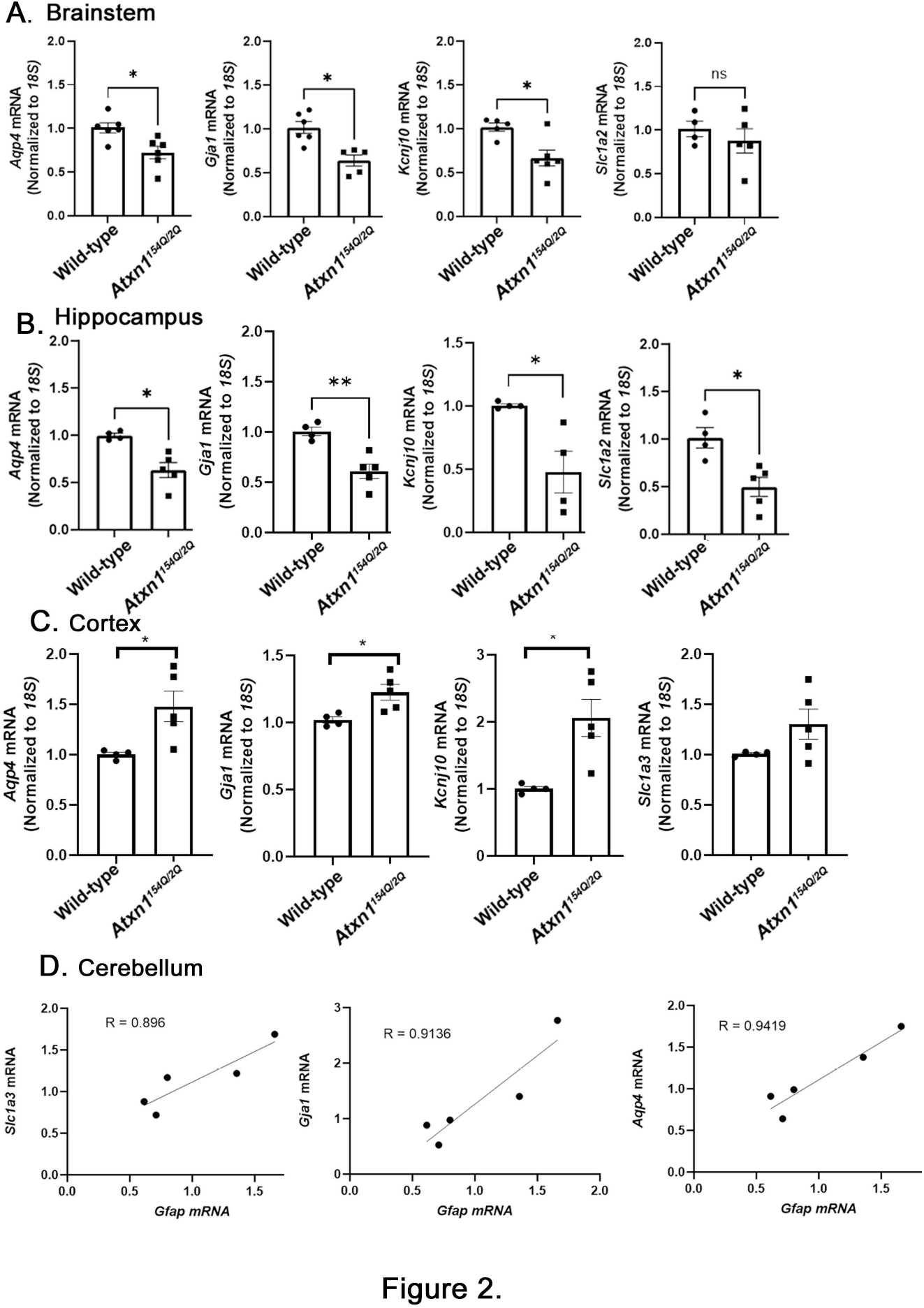
Spatial diversity of molecular changes in astrocytes during early SCA1. mRNA was extracted from the brainstem (A, medulla), hippocampus (B), cortex (C), and cerebellum (D) of 12 weeks old *Atxn1*^*154Q/2Q*^ and wild-type littermate controls (N = 4-6) and RTqPCR was used to evaluate expression of astrocyte specific genes. (A-C) Data is presented as mean ± SEM with average values for each mouse represented by a dot. * p<0.05 Students t-test. (D) Cerebellum RTqPCR data is presented as Pearson’s correlation between *Gfap* and homeostatic astrocyte genes.

In the cerebellum, we have found a large variability in the expression of *Aqp4, Gja1, Kcnj10*, and *Slc1a3*, that may reflect large variability in the reactive state of cerebellar astrocytes (as determined by the *Gfap* expression). Indeed, we found a positive correlation between the expression of *Aqp4, Gja1*, and *Slc1a3*, and the expression of *Gfap* (Figure 2D), and a trending correlation between *Gfap* and *Kcnj10*. These results indicate that early in disease, cerebellar astrocytes with a higher *Gfap* expression (denoting that they are likely to be more reactive) also have a higher expression of homeostatic genes

### Regional diversity of microglial SCA1 changes

We have previously shown that microglia are reactive in SCA1 cerebellar cortex and contribute to SCA1 pathogenesis in different SCA1 mouse models [65][66]. We investigated changes in microglial density and morphology in the cerebellum and other brain regions using IHC staining for the microglial marker ionized calcium binding adaptor protein (Iba1). We quantified microglial density and the percent area covered by microglial Iba1+ processes (Figure 3). In the cerebellum, we found a significant increase in microglia density and Iba1+ percent area in 12 week old *Atxn1*^*154Q/2Q*^ mice (Figure 3A). Similarly, in the brainstem (Figure 3C) and cortex (Figure 3D) we found an increase in microglial density. Microglial area, indicative of hypertrophy, was significantly increased only in the cortex. We found no significant changes in microglial density nor percent area in the hippocampus (Figure 3B).

**Figure 3.**
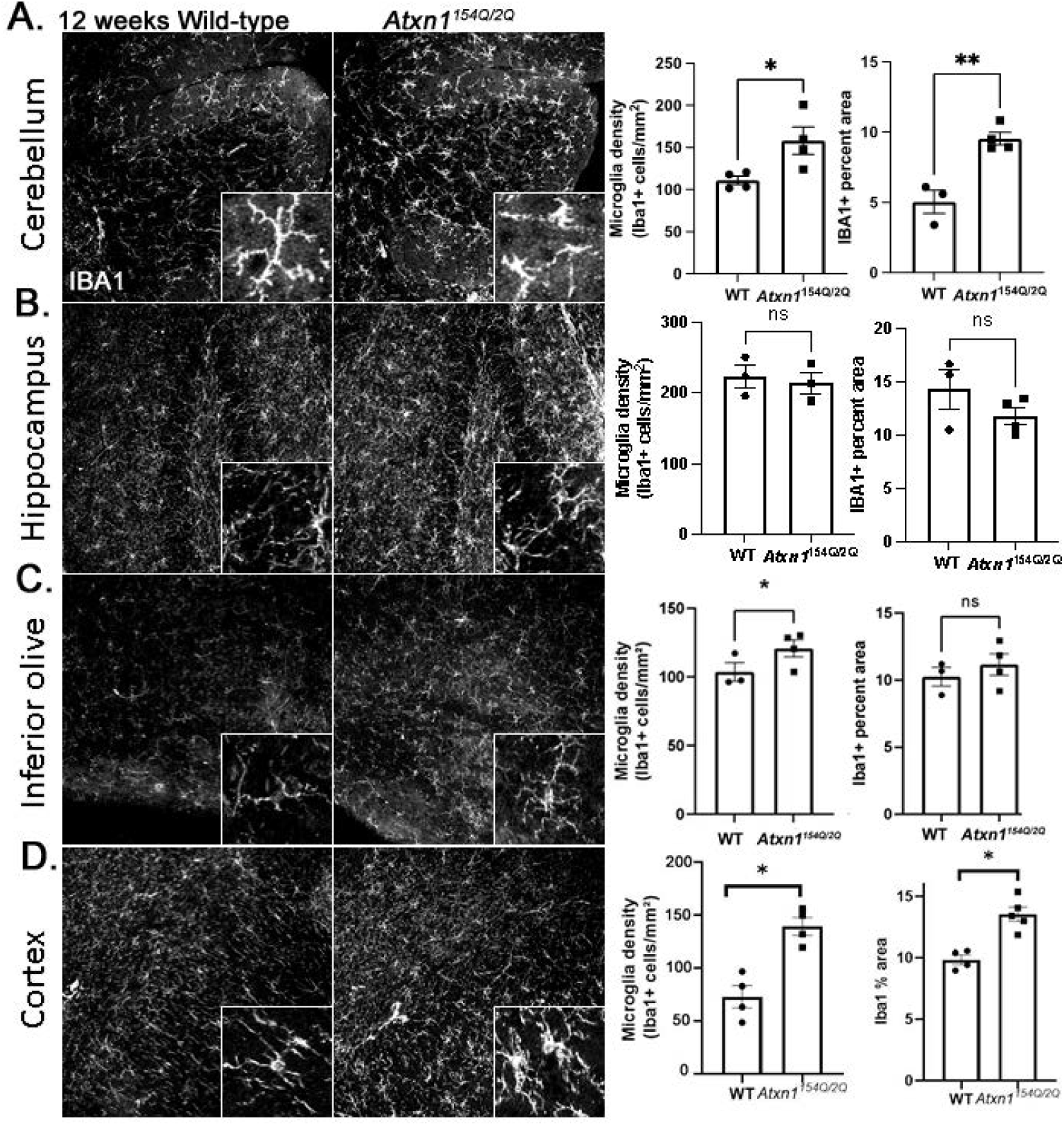
Early SCA1 microglial morphology in *Atxn1*^*154Q/2Q*^ mice. Brain sections from 12 weeks old *Atxn1*^*154Q/2Q*^ and wild-type littermate controls (N = 3-5) were stained for Iba1. Confocal images of cerebellum (A, lobule X), hippocampus (B, dentate gyrus), brainstem (C, olivary nucleus), and motor cortex (layer 6 and underlying corpus callosum) were used to quantify density of Iba1 positive microglia and Iba1 area. Data is presented as mean ± SEM with average values for each mouse represented by a dot. * p<0.05 Students t-test.

Reactive changes in glia can be caused by neuronal loss. To determine whether observed glial changes may be caused my neuronal loss we used neuronal marker NeuN to quantify neuronal numbers. We did not find evidence of neuronal loss in any of the examined brain regions at this early disease stage in *Atxn1*^*154Q/2Q*^ mice (Supplementary Figure 1). While this result does not exclude neuronal dysfunction, our results suggest that spatially diverse glial phenotypes precede neuronal loss.

To investigate whether changes in the expression of core astrocytic genes may be associated with more subtle neuronal changes, we quantified neuronal activity in hippocampus and cortex, two regions with reduced and increased expression of core astrocyte genes. Intriguingly, ratio active cFos+ neurons (cFos+/NeuN+) was decreased in hippocampus while it was not changed in motor cortex of *Atxn1*^*154Q/2Q*^ mice (Supplementary Figure 2A-B).

Together these results suggest that microglia—similar to astrocytes—exhibit spatial diversity of SCA1 changes. Intriguingly, while early in disease astrocytes undergo both loss and hypertrophy, microglia mostly undergo hypertrophy, indicating a complex orchestration of astrocyte and microglial phenotypes early in SCA1. Moreover, decreased astrocytic support may contribute to reduced neuronal activity in hippocampus.

### Temporal diversity of astrocyte phenotypes in SCA1s

We next investigated how glial phenotypes changes with disease progression. At late stage of disease (20 weeks) astrocytes in the cerebellum, hippocampus and motor cortex showed signs of hypertrophy (Figure 4 A, B, D). In olives the loss and hypertrophy of astrocytes were more pronounced (Figure 4 C).

**Figure 4.**
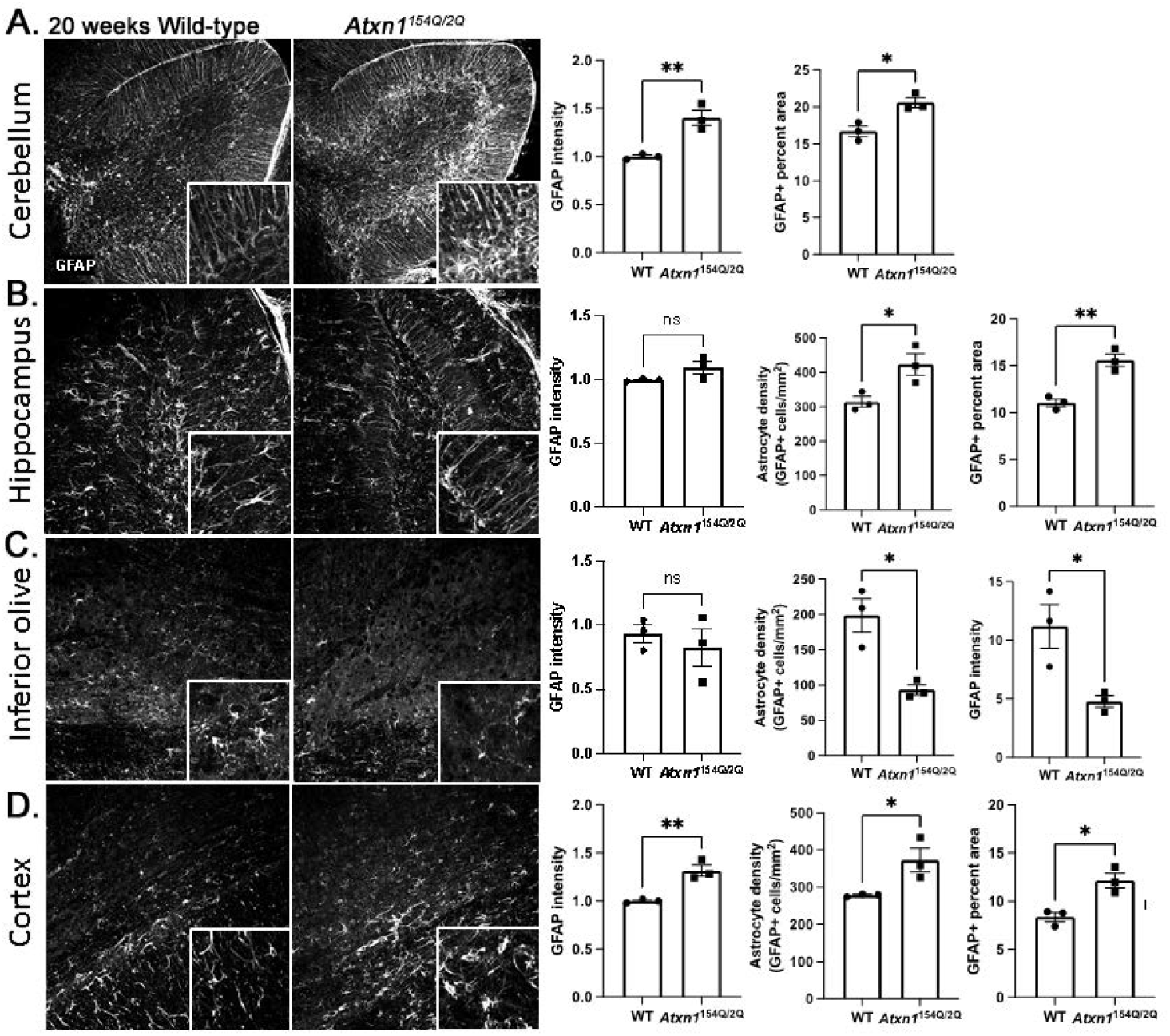
Spatial diversity of astrocytic morphology in *Atxn1*^*154Q/2Q*^ mice at late disease stage. Brain sections from 20 weeks old *Atxn1*^*154Q/2Q*^ and wild-type littermate controls (N = 3-5) were stained for GFAP. Confocal images of cerebellum (A, lobule X), hippocampus (B, dentate gyrus), brainstem (C, olivary nucleus), and motor cortex (layer 6 and corpus callosum) were used to quantify GFAP intensity, density of astrocytes and GFAP area. Data is presented as mean ± SEM with average values for each mouse represented by a dot. * p<0.05 Students t-test.

Using RNA sequencing we compared the expression of astrocytic genes in these regions of *Atxn1*^*154Q/2Q*^ mice and their wild-type littermate controls. We found a significant decrease in the expression of astrocytic homeostatic genes *Kcnj10, Aqp4, Gja1, and Slc1a2* in all four regions in the late stage *Atxn1*^*154Q/2Q*^ mice (Figure 5). These results indicate temporal diversity of astrocyte phenotypes in SCA1 and suggest that brain-wide loss of astrocyte core gene expression characterizes late stages of disease in *Atxn1*^*154Q/2Q*^ mice.

**Figure 5.**
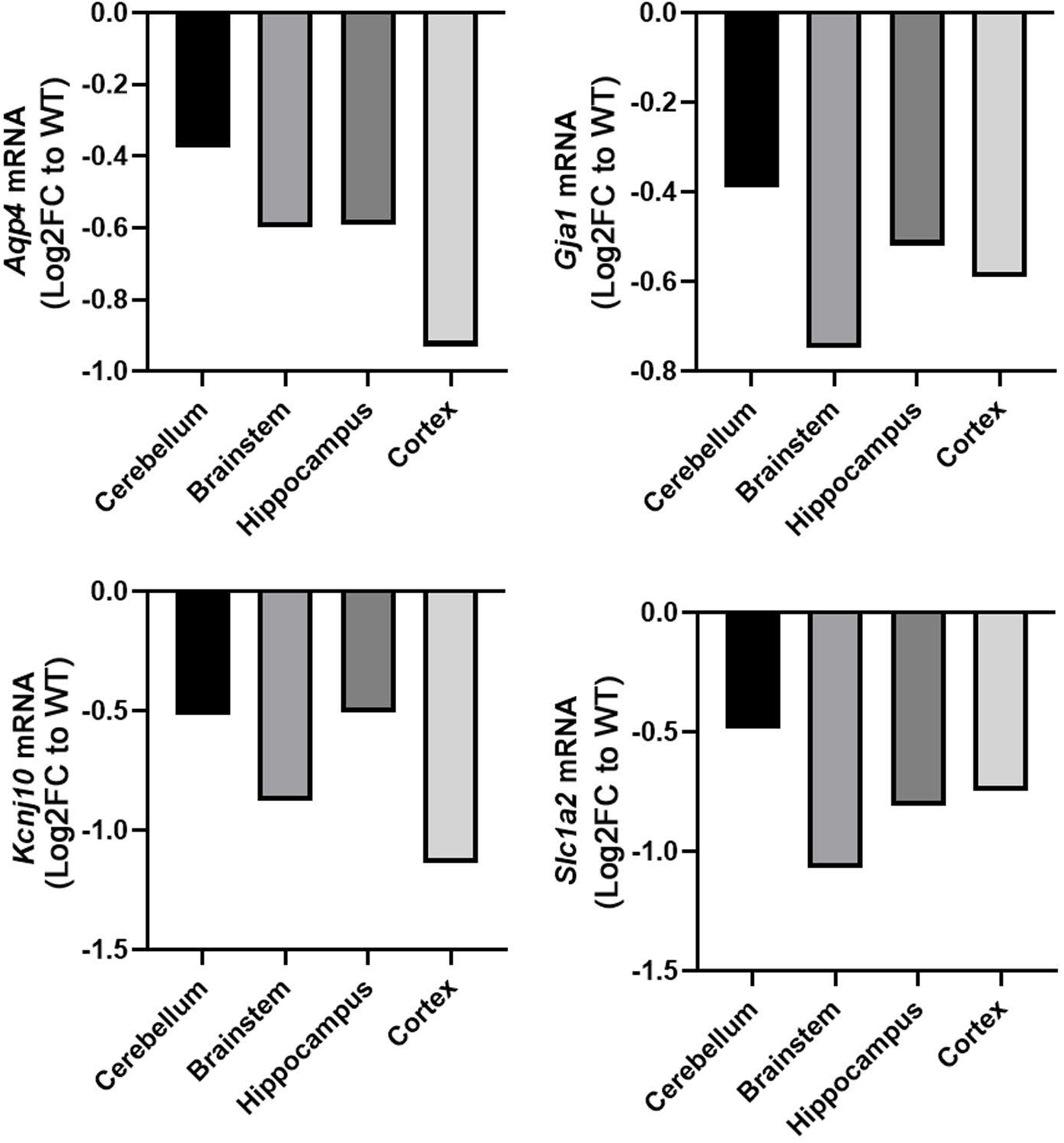
Late SCA1 stage astrocyte molecular analysis indicates brain wide loss of core homeostatic functions. Cerebellum, brainstem (medulla), hippocampus, and cortex were isolated from 26 weeks old *Atxn1*^*154Q/2Q*^ and wild-type control mice (N=4 of each genotype) and isolated RNA was sent for RNA-sequencing (RNA-seq). Expression of astrocyte genes was determined using differential gene expression analysis with numbers in each square representing log2FC in *Atxn1*^*154Q/2Q*^ compared to wild-type controls. All presented results have p<0.05.

Intriguingly, late in diseases both olives and hippocampus demonstrated decrease in microglial density (Figure 6). While density of cerebellar microglia was further increased with disease progression, we did not detect any change in microglial density in the cortex. As microglia have been suggested to lead to loss of synapses in disease we quantified excitatory synapses in the cerebellum, that exhibit microglial activation and cortex and hippocampus, that show reduction or no changes in microglial density. We found significant reduction of VGLUT2 synapses on the Purkinje cells in the cerebellum, while there was no change in the VGLUT2/PSD95 synaptic quanta in hippocampus and motor cortex (Supplementary Figure 3).

**Figure 6.**
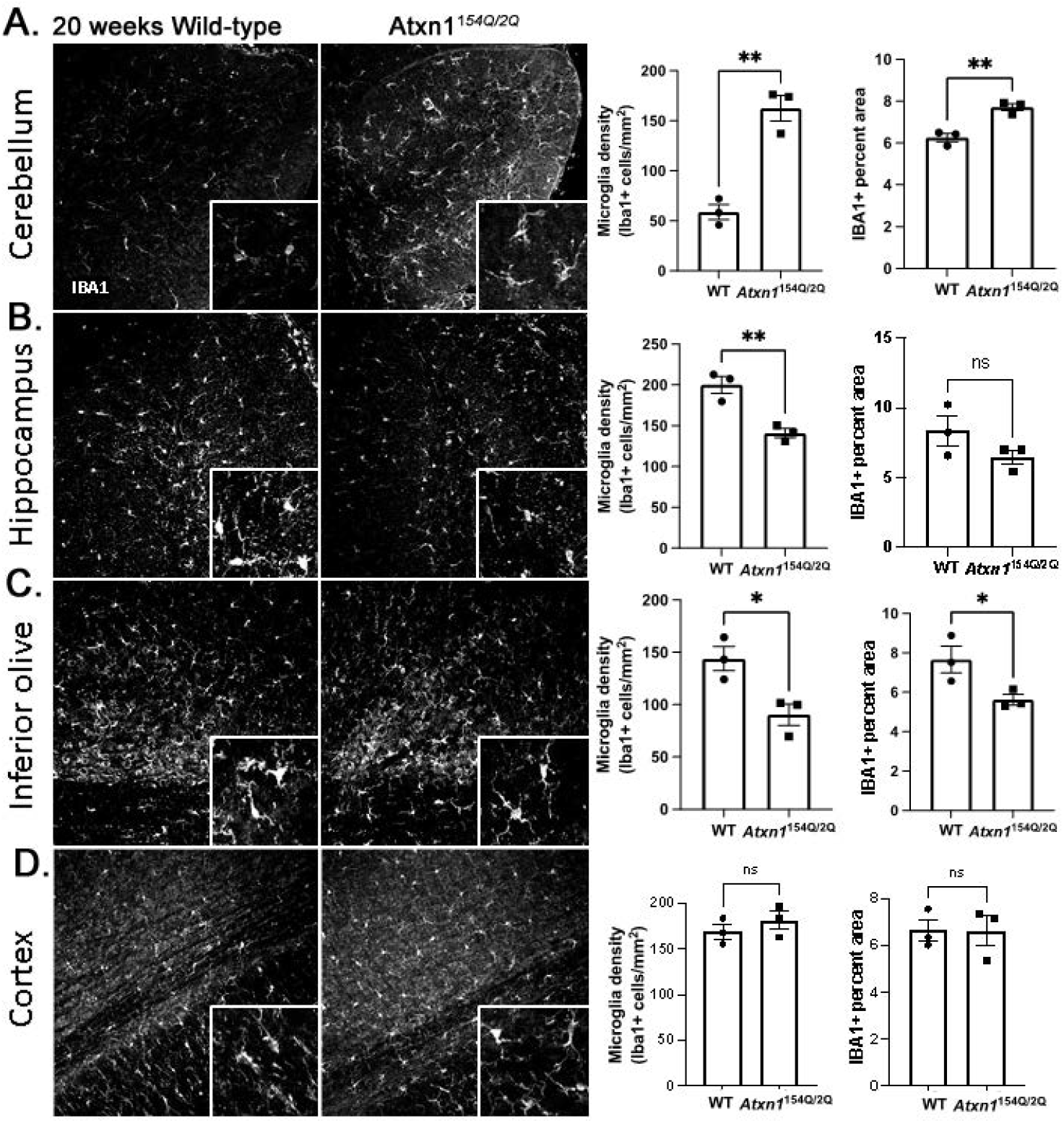
Diversity of late SCA1 microglial morphology phenotypes in *Atxn1*^*154Q/2Q*^ mice. Brain sections from 20 weeks old *Atxn1*^*154Q/2Q*^ and wild-type littermate controls (N = 3-5) were stained for Iba1. Confocal images of cerebellum (A, lobule X), hippocampus (B, dentate gyrus), brainstem (C, olivary nucleus), and motor cortex (layer 6 and corpus callosum) were used to quantify density of Iba1 positive microglia and Iba1 area. Data is presented as mean ± SEM with average values for each mouse represented by a dot. * p<0.05 Students t-test.

These results suggest that with disease progression some brain regions, such as olives may suffer from loss of both astrocytes and microglia, while other regions, such as cerebellum may suffer from reactive phenotypes in both cell types.

## 4. Discussion

Here, we report on the spectrum of reactive astrocyte phenotypes, across several clinically relevant brain regions of *Atxn1*^*154Q/2Q*^ mice and at different stages of disease progression. Recent studies indicated heterogeneity of glial cells across healthy brain regions [16] and across different disease conditions [67][56]. Our results indicate spatial diversity of morphological and molecular changes in astrocytes during the early stages of disease in *Atxn1*^*154Q/2Q*^ mice. This spectrum of reactive astrocytic phenotypes may indicate brain-region-specific astrocytic dysfunctions and consequently their brain region specific contributions to SCA1 disease pathogenesis. Given that these changes precede neuronal loss, it is crucial to examine glial heterogeneity as a means to distinguish protective versus deleterious astrocytic phenotypes and identify opportunities for early therapeutic intervention. For this reason, we focused here of the astrocytic neuroprotective genes *Slc1a2, Kcnj10, Gja1*, and *Aqp4*, and found that their expression was reduced in the medulla and hippocampus, but was increased in the cortex of *Atxn1*^*154Q/2Q*^ mice at 12 weeks of age and before notable neuronal loss [6]. *Kcnj10* encodes a potassium rectifier, Kir4.1, that is involved in maintaining potassium homeostasis [68] and has been implicated in several diseases, including Huntington’s disease, ALS, and depression [69][68][70]. Gja1 encodes the astrocyte gap-junction protein connexin 43 (*Gja1)* which is critical for astrocyte-astrocyte communication [50]. Aquaporin (*Aqp4)*, is critical for both interstitial fluid drainage in the glymphatic system and water homeostasis [71]. Astrocytic *Slc1a* (2 and 3) genes encode glutamate transporters that are responsible for removing glutamate from synaptic space. As such, these astrocytic genes are critical for neuronal function, and have been implicated in the pathogenesis of several neurodegenerative diseases [72][73]. We propose that reduced expression of astrocyte neuro-supportive genes—as observed in *Atxn1*^*154Q/2Q*^ mice— may exacerbate neuronal dysfunction and SCA1 pathogenesis in the hippocampus and medulla. Conversely, increased expression of these neuro-supportive astrocyte genes in the cortex may indicate compensatory roles of astrocytes that delay neuronal dysfunction in the SCA1 affected cortex. This is supported by our results showing preserved neuronal activity in the cortex and reduced neuronal activity in the hippocampus.

Early in disease, we found no evidence of increased GFAP expression, a hallmark of reactive astrogliosis, in the medulla. Likewise, we found only trending indications of increased GFAP expression in the cortex. Thus, our results may indicate that increased GFAP expression may not negatively correlate to changes in astrocyte homeostatic gene expression in disease conditions as previously suggested. This is consistent with a previous study which demonstrated that reduced expression of the astrocyte homeostatic gene *Kcnj10* in the striatum of Huntington’s mice is not dependent on astrogliosis as defined as increased GFAP expression [68]. Moreover, in our study we have found a strong positive correlation of *Gfap* expression with the expression of *Kcnj10, Slc1a2* and *Aqp4 i*n the cerebellum, indicating that reactive astrocytes (indicated by increased Gfap) may initially increase expression of core homeostatic genes as a neurosupportive measure in disease.

We also found a loss of astrocytes in the medulla of 12-week-old *Atxn1*^*154Q/2Q*^ mice, preceding neuronal loss. This is consistent with previous work showing lower levels of *myo*-inositol (Ins), an astrocytic marker, in the brainstem of 18-and 28-week-old *Atxn1*^*154Q/2Q*^ mice [33]. Moreover, treatment with *Atxn1* targeting antisense oligonucleotides (ASO), which are also taken up by astrocytes [74], was shown to rescue lower Ins levels in the brainstem and prolong the survival of *Atxn1*^*154Q/2Q*^ mice [33]. Based on these results we propose that reduced expression of homeostatic genes and decreased astrocyte viability may contribute to brainstem dysfunction and premature lethality in SCA1 mice. However, while neuronal numbers were not altered in medulla of 12-week-old *Atxn1*^*154Q/2Q*^ mice, we cannot exclude neuronal dysfunction. Future studies using conditional SCA1 mice to selectively delete expression of mutant ATXN1 in astrocytes will provide insight into the cell autonomous effect of mutant ATXN1 on astrocyte pathology in the medulla and other brain regions in SCA1.

Finally, we report that disease progression eliminates diversity of astrocyte gene expression changes across brain regions. On the other hand morphological changes in astrocytes became more pronounced with disease. As disease progressed hippocampus and brainstem exhibited decrease in microglial density. Together, these results indicate that profound loss of glial functions characterizes late disease stage in *Atxn1*^*154Q/2Q*^ mice and is likely to exacerbate neuronal dysfunction and loss at this stage of disease (Figure 7). While future studies will investigate mechanisms underlying the loss of astrocyte and microglia, our results indicate that preserving glial functionality may provide therapeutic benefits in SCA1.

**Figure 7.**
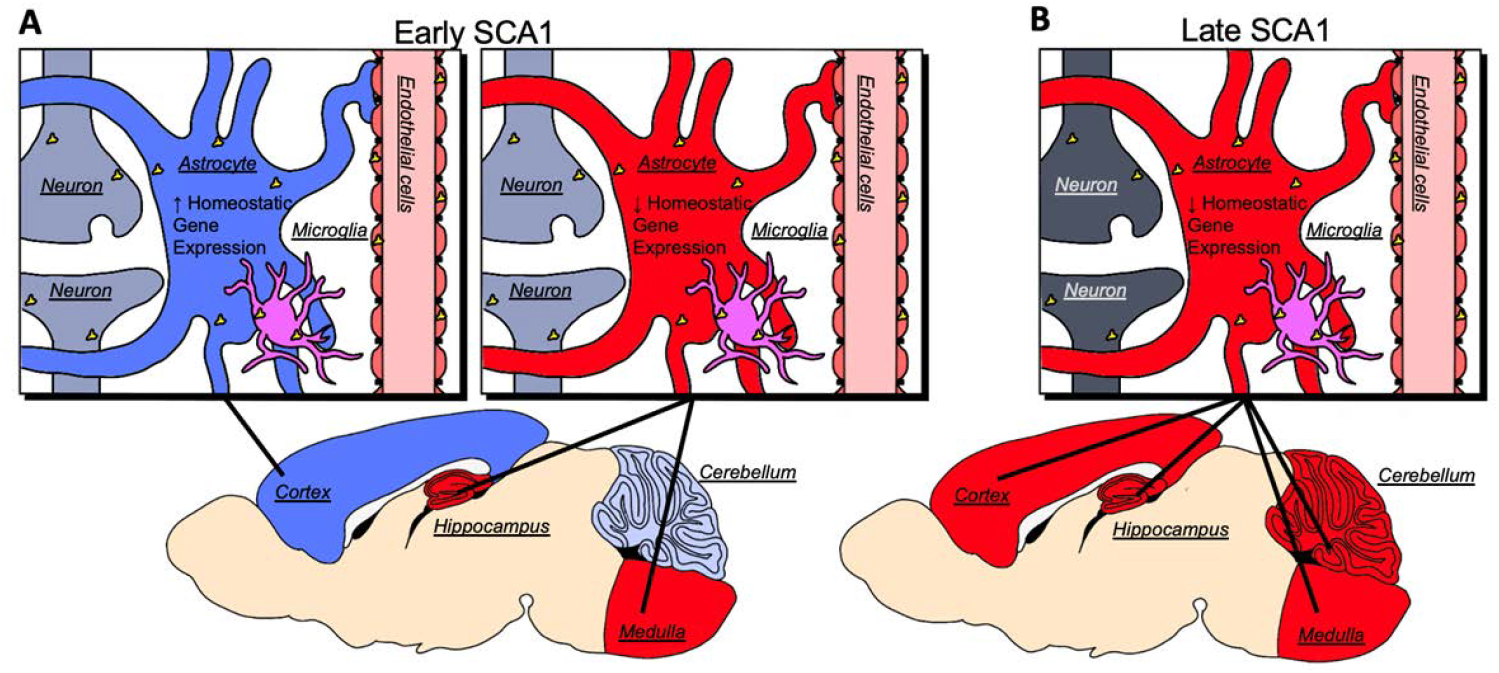
Simplified schematics of astrocytic changes in SCA1. (A) Early stages of SCA1. Brain regions indicated in blue exhibit an increase (cortex) or slight change (cerebellum) in the expression of astrocyte homeostatic genes whereas regions in red exhibit loss of astrocyte homeostatic genes (hippocampus and medulla oblongata). (B) Late, terminal stages of SCA1. Astrocytes in all brain regions exhibit reduced expression of core genes necessary for astrocyte homeostatic function (cortex, cerebellum, hippocampus, and medulla). Microglia activation in different brain regions does not correlate with astrocyte reactivity or homeostatic gene expression. Yellow represents cell-types shown to express the *Atxn1* gene in mice and ATXN1 humans.

## Acknowledgements

We acknowledge all the members of Orr and Cvetanovic laboratories for thoughtful discussions and feedback on the study. Work in this study was aided by the Mouse Behavioral Core and Genomics Core at the University of Minnesota. We thank Laura Berg for editing.

## Data Availability

Data available on request from the authors.

## Author Contributions

M.C. conceptualized the study. J-G.R., C.S., K.H., A.S., and S.G. performed experiments. J-G.R., M.C., F.G., and C.S. analyzed data. M.C., and J-G.R., wrote the manuscript with input from H.T.O., A. K., C.S., K.H., A.S., and S.G.

## Supplementary Figures

**Supplementary Figure 1. Neuronal density is not altered early in SCA1**. Brain sections from 12 weeks old *Atxn1*^*154Q/2Q*^ and wild-type littermate controls (N = 3-5) were stained for NeuN. Confocal images of cerebellum (A, deep cerebellar nuclei), hippocampus (B, CA2/CA3), brainstem (C, olivary nucleus), and motor cortex (layer 6) were used to quantify density of NeuN+ neurons. Data is presented as mean ± SEM with average values for each mouse represented by a dot. * p<0.05 Students t-test.

**Supplementary Figure 2. Neuronal activity is altered in the hippocampus but preserved in motor cortex early in SCA1**. Brain sections from 12 weeks old *Atxn1*^*154Q/2Q*^ and wild-type littermate controls (N = 3-5) were stained for NeuN and c-Fos. Confocal images of hippocampus (A, dentate gurys), and motor cortex (B, layer 6) were used to quantify density of NeuN+ and c-Fos+ neurons. Data is presented as a mean ratio of c-Fos/NeuN+ cells ± SEM with average values for each mouse represented by a dot. * p<0.05 Students t-test.

**Supplementary Figure 3. Synaptic quantification in the cerebellum, hippocampus and cortex**. Brain sections from 18 weeks old *Atxn1*^*154Q/2Q*^ and wild-type littermate controls (N = 3-5) were stained for VGLUT2, and calbindin (A, cerebellum) or VGUT2 and PSD95 (B, hippocampus and cortex). Confocal images were used to quantify ratio of length of VGLUT2 puncta over calbindin+ PC dendrites (A, cerebellum), or number of VGLUT2 and PSD 95 puncta and their overlay (B, hippocampus and cortex). Data is presented as a mean ± SEM with average values for each mouse represented by a dot. * p<0.05 Students t-test.

