## Supplementary figures and images for "Spatial and temporal diversity of astrocyte phenotypes in Spinocerebellar ataxia type 1 mice"

### Supplementary Figures

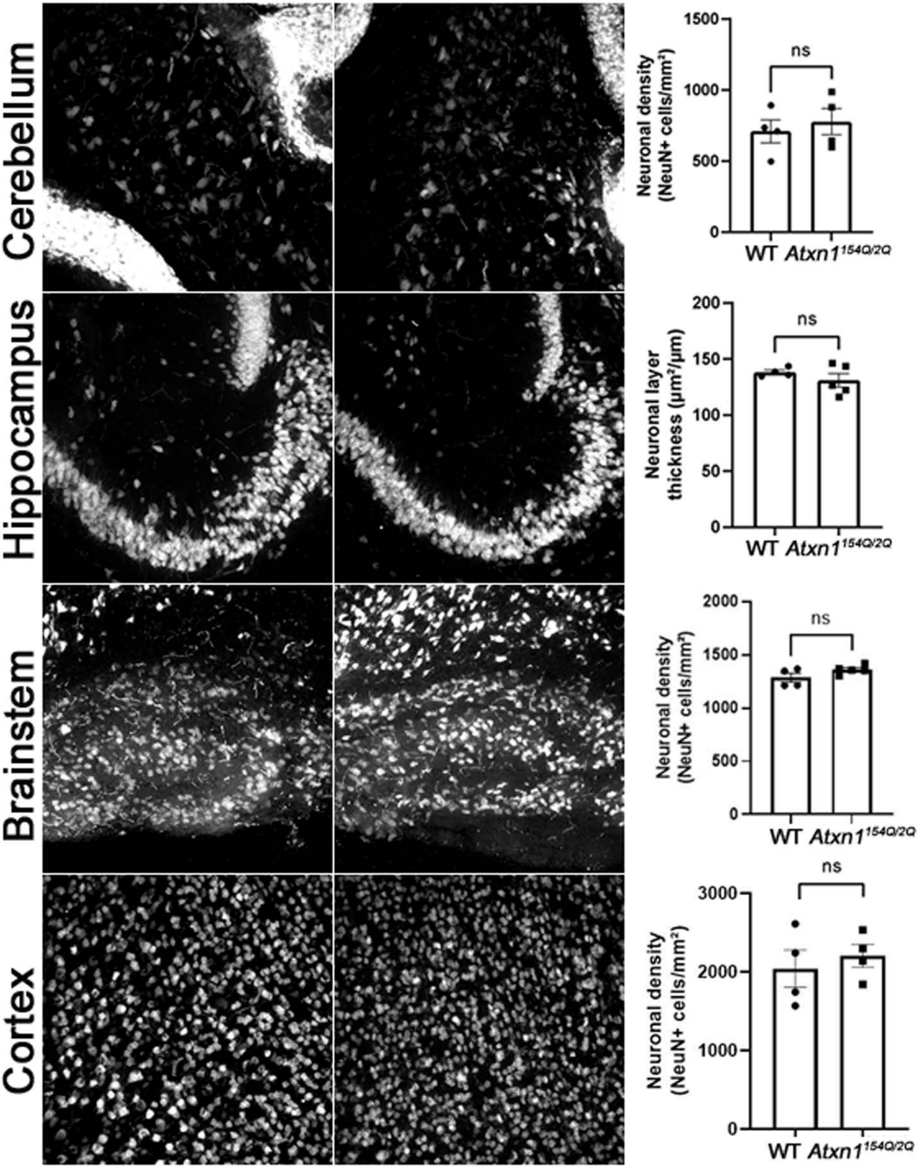


#### Supplementary Figure 2.

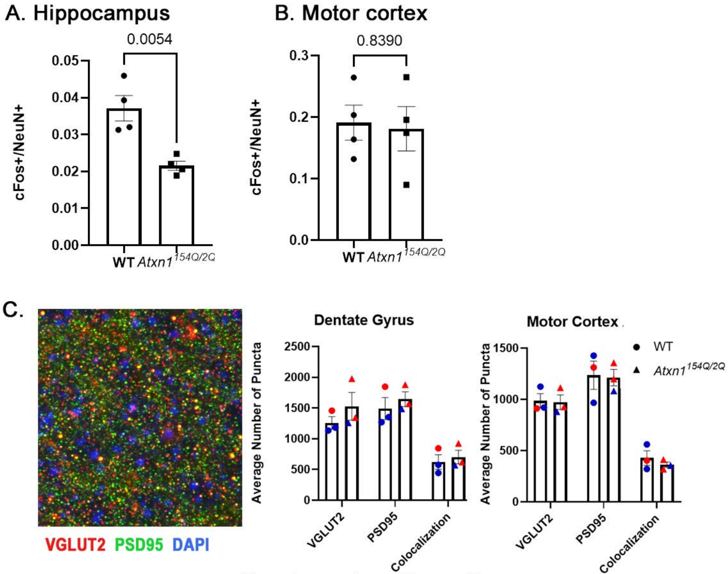


Supplementary Figure 2.

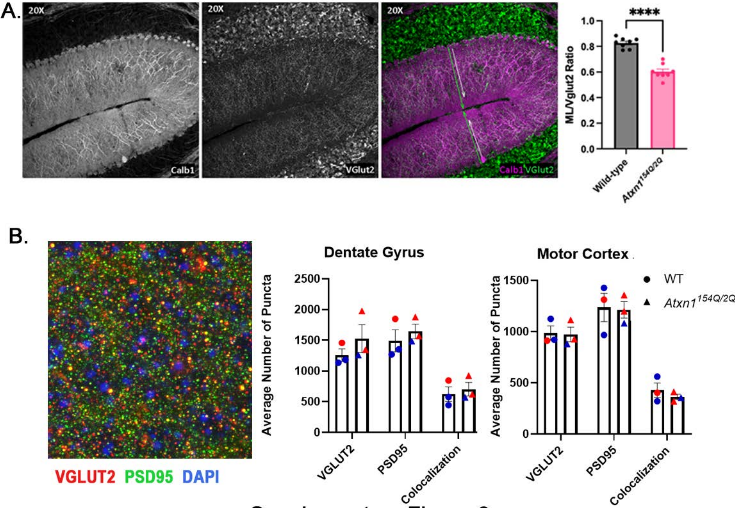


Supplementary Figure 3.
